# HIV-1 Integrase C-terminal Domain Mutations Regulate ALLINI Sensitivity

**DOI:** 10.64898/2026.10.22.761050

**Authors:** Abhishek Kumar, Tung Dinh, Christian Shema Mugisha, Anthony Boateng, Qianzi Zhou, Mamuka Kvaratskhelia, Sebla B. Kutluay

**Affiliations:** Department of Molecular Microbiology, Washington University School of Medicine, Saint Louis, Missouri, USA; Division of Infectious Diseases, University of Colorado School of Medicine, Aurora, Colorado, USA

## Abstract

HIV-1 integrase (IN) performs two essential functions in virus replication: catalyzing the integration of reverse-transcribed viral cDNA into host DNA and promoting proper virion maturation. The latter is mediated by IN binding to the genomic RNA (gRNA) through basic residues in its C-terminal domain (CTD), thereby ensuring correct repackaging of the gRNA inside the remodeled capsid lattice. Allosteric integrase inhibitors (ALLINIs) bind within a V-shaped pocket at the interface of the catalytic core domain (CCD) of IN dimers, promote the engagement of the CTDs from adjacent dimers, and thereby occlude gRNA binding. However, virological evidence supporting a role for the IN-CTD in ALLINI mechanism of action is lacking, as mutations in CTD residues implicated in ALLINI-induced multimerization typically yield non-infectious viruses. Here we demonstrate that IN-CTD mutants bearing compensatory substitutions that restore infectivity exhibit markedly altered sensitivity to ALLINIs. Specifically, we show that R269A/K273A substitutions within the IN-CTD confer ∼100-fold enhanced sensitivity to ALLINIs. This correlates with a loss of IN-gRNA binding at low ALLINI concentrations, while exerting minimal effects on IN multimerization *in vitro* and in virions. In contrast, we found that the IN I268T substitution which arose as a suppressor mutation of I268A confers partial resistance to ALLINI-induced inhibition of gRNA binding. Collectively, these findings provide the first direct genetic evidence that CTD residues, despite being distal to the primary drug binding site at the CCD dimer, modulate sensitivity to ALLINIs and highlights IN-gRNA binding as a central target of ALLINIs.

**SIGNIFICANCE:** In addition to its catalytic role in viral DNA integration, HIV-1 integrase (IN) is essential for virion maturation. IN binding to the viral RNA genome (gRNA) through basic residues in its C-terminal domain (CTD) ensures the proper repackaging of the gRNA inside the mature capsid lattice. Allosteric integrase inhibitors (ALLINIs) bind to a pocket within the catalytic core domain (CCD) of an IN dimer and are thought to disrupt IN-gRNA interactions by triggering aberrant IN multimerization through the engagement of the CTDs of a nearby dimer. However, direct genetic evidence supporting a role for the IN-CTD in ALLINI mechanism of action is lacking. Herein, we show that IN-CTD mutants bearing compensatory substitutions that restore infectivity have significantly altered sensitivities against ALLINIs. Our findings provide the first virological evidence that residues within the CTD, away from the primary drug binding site at the CCD dimer, can alter ALLINI sensitivity.

## INTRODUCTION

HIV-1 integrase (IN) performs two essential functions during virus replication. First is its well-defined function in covalent insertion of reverse-transcribed viral DNA (vDNA) into target cell chromatin (1, 2). Second, IN plays a non-catalytic role during virion maturation by binding to the viral genomic RNA (gRNA), thereby ensuring proper repackaging of the viral ribonucleoprotein (vRNP) complexes within the remodeled mature capsid lattice (3). The catalytic function of IN has been successfully targeted by integrase strand-transfer inhibitors (INSTIs), including raltegravir, dolutegravir, bictegravir and cabotegravir (4–9). Although INSTIs are central components of anti-retroviral therapy regimens, the emergence of drug-resistant HIV-1 variants remains a persistent challenge (10–14). Targeting the non-catalytic function of IN through allosteric IN inhibitors (ALLINIs), a novel class of investigational antiretrovirals, offers an alternative therapeutic strategy that can complement INSTI-based regimens.

Structurally, IN comprises three independently folded domains: the N-terminal domain coordinates Zn^2+^ via a conserved His-His-Cys-Cys (HHCC) motif; the catalytic core domain (CCD) adopts an RNase H superfamily fold and contains a D, D-35-E motif, responsible for DNA cleavage and strand transfer; and the CTD contains an Src homology domain 3 (SH3)-like fold that contributes to DNA binding (15, 16). Higher order oligomerization of IN is required for its catalytic activity (17) as supported by recent cryogenic electron microscopy (cryo-EM) studies of HIV-1 intasomes indicating that integration is likely mediated by dodecameric assemblies (18, 19).

Another important function of IN is the regulation of proper virion maturation (20). This non-catalytic role was initially uncovered through extensive mutational analyses of IN (21). While a subset of mutations predictably inhibited the catalytic function of IN (21–23), a second subset, collectively referred to as class II mutations, exhibited pleiotropic effects on virus replication, including defects in particle assembly (22, 24–36), morphogenesis (20, 22, 26, 32–34, 37, 38), and reverse transcription in target cells (20–22, 28, 30–32, 34, 36–54). Notably, several of these mutations did not impair IN catalytic function in vitro (26, 27, 30, 31, 40, 41, 44, 46, 55–57), emphasizing that the catalytic and non-catalytic functions of IN are separable. A hallmark of class II IN mutant viruses is the mislocalization of vRNPs, which are found eccentrically positioned outside of the viral capsid (CA) lattice (20, 22, 26, 32–34, 37, 38). Subsequent studies demonstrated that IN binding to the gRNA is critical for virion maturation and underlies the pleiotropic defects observed in many class II IN mutants (20, 58).

A critical determinant of IN binding to gRNA is proper IN oligomerization. IN tetramers exhibit substantially higher affinity towards RNA than IN monomers and dimers (58) and recent cryo-EM studies indicate that a linear filament made of IN octamer repeat units bind RNA and engage with the CA lattice (59). While basic residues within the IN-CTD (e.g. R262, R263, R269, K273) directly bind to the gRNA through electrostatic interactions (20, 58–60), class II IN mutations located outside the CTD can impair IN tetramerization and indirectly inhibit IN-gRNA binding in virions (58). ALLINIs induce aberrant IN multimerization (61–67) by binding to a V-shaped pocket at the dimer interface formed by the CCDs and subsequently recruiting CTDs from adjacent dimers (68, 69). Consequently, ALLINI-induced aberrant hyper-multimerization of IN is thought to disrupt functional tetramer formation and preclude IN-gRNA interactions.

Cell culture based viral breakthrough studies in the presence of ALLINIs have identified multiple drug-resistant substitutions near the inhibitor binding site at the CCD dimer interface (e.g. Y99H, A128T, H171Q, T174I). By contrast, no resistance mutations have been detected in the CTD. It should be noted that the CCD contains several variable amino acids near the inhibitor binding site, whereas the CTD residues interacting with the CCD + ALLINI complex are either invariant or highly conserved (68, 69). While substitutions of the ALLINI-interacting CTD residues in the context of full-length recombinant IN confer marked resistance to ALLINI-induced aberrant IN multimerization (68, 69), the same CTD amino acid mutations in the context of HIV-1 result in non-infectious particles, thereby hampering efforts to delineate the contribution of the CTD to the antiviral activity of ALLINIs in relevant biological settings.

Herein we aimed to gain direct virological evidence for the proposed involvement of IN-CTD in the ALLINI mechanism of action. In agreement with prior findings, we found that Ala substitution of Y226, W235, K266, I268 residues within the IN-CTD implicated in ALLINI-mediated multimerization resulted in a class II phenotype and yielded non-infectious virions in which IN-gRNA interactions were inhibited. To overcome this limitation, we examined the ALLINI sensitivities of previously described class II IN-CTD mutants (e.g. R262A/R263A and R269A/K273A) bearing compensatory charge reversal substitutions at nearby residues (e.g. D256R, D278R) that restore infectivity (60). We found that compared to WT HIV-1, HIV- 1_(R269A/K273A/D278R IN)_ was ∼100-fold more sensitive to quinoline-based ALLINIs, BI-D and BI-B2, as well as pyrrolopyridine-based Pirmitegravir. The enhanced sensitivity of HIV-1_(R269A/K273A/D278R IN)_ to ALLINIs mapped to the deleterious R269A/K273A mutations but not the compensatory D278R substitution and correlated with loss of gRNA binding but not enhanced aberrant IN multimerization. In addition, through serial passaging experiments, we isolated I268V and I268T/D270N substitutions as suppressors for the IN I268A mutant. IN I268T/D270N was markedly less sensitive to ALLINI-induced loss of gRNA binding whereas the more conserved IN I268V mutant displayed similar sensitivity as WT IN. Taken together, our findings provide the first virological evidence that residues within the CTD, distal from the primary drug binding site at the CCD dimer, can alter ALLINI sensitivity.

## RESULTS

### Mutations of the CTD residues involved in ALLINI-induced aberrant IN multimerization display class II phenotypes

Structural and biochemical studies have shown that CTD residues Y226, W235, K266 and I268 mediate binding to the ALLINI-bound CCD (**Fig. 1A**) and alanine (Ala) substitutions at these residues inhibit ALLINI-induced aggregation *in vitro* (68, 69). We first examined the properties of viruses bearing Ala substitutions at these residues within the context of full-length HIV-1 NL4-3 molecular clone. Y226A, W235A, K266A and I268A substitutions did not affect Gag/Gag-Pol expression or processing in cells (**Fig. 1B**). Similar levels of particles were released in the culture supernatants, though we noted a modest reduction in IN incorporation into virus particles (**Fig. 1B**). The resulting viruses exhibited markedly reduced infectivity (**Fig. 1C**) and displayed a class II phenotype characterized by substantially lower levels of IN-RNA complexes isolated from virions (**Fig. 1D, 1E**). We noted that although similar amounts of IN were incorporated in virus particles, the Y226A, W235A and I268A mutants yielded considerably lower amount of IP’ed IN protein, suggesting that the introduced mutations destabilize IN during the IP procedure (**Fig. 1D, 1E**). In contrast, IN K266A substitution yielded comparable levels of IP’ed IN protein as WT but was nonetheless impaired in RNA binding (**Fig. 1D, 1E**).

**Figure 1.**
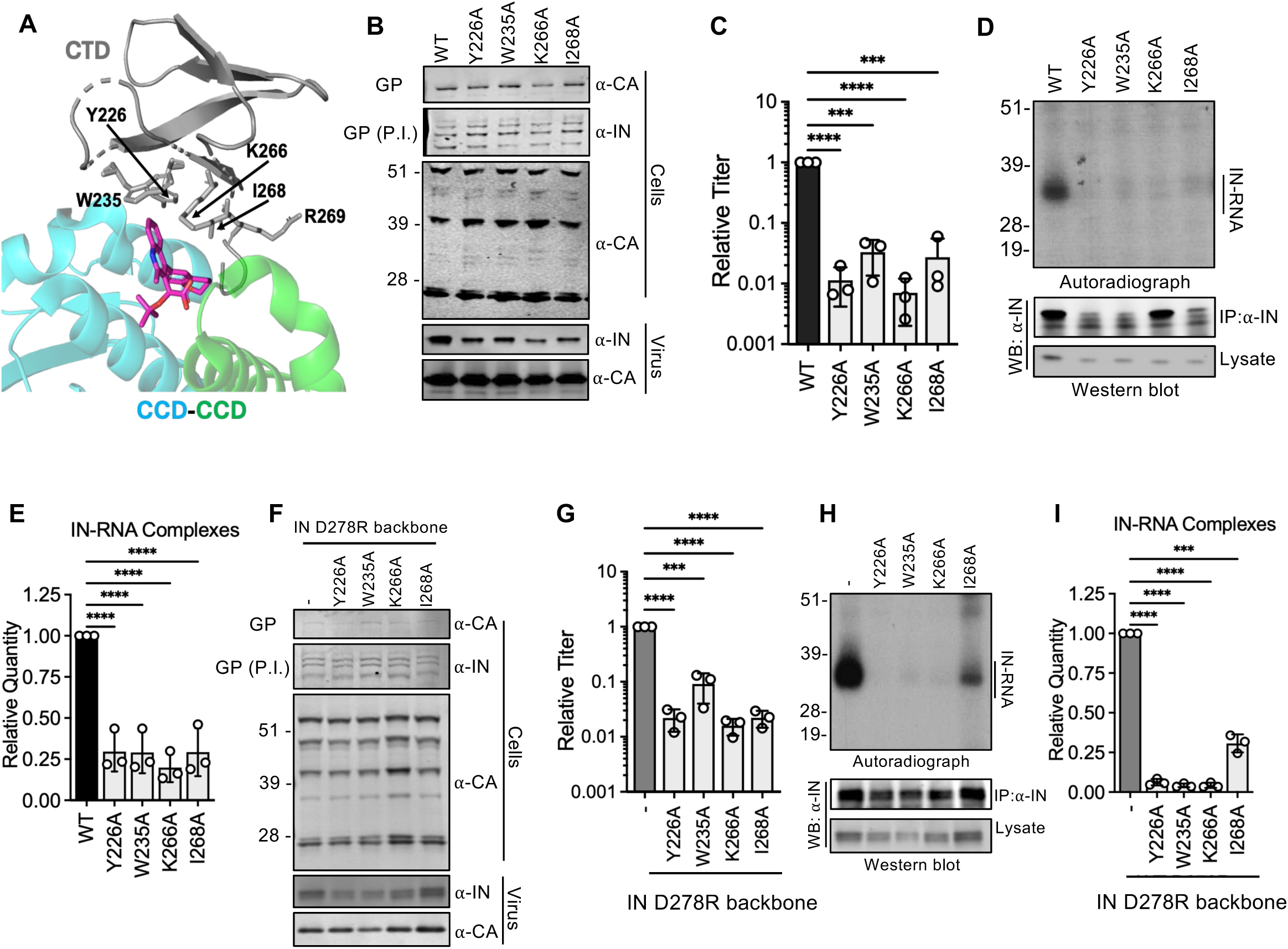
Mutations in ALLINI-interacting CTD residues result in non-infectious viruses with class II phenotypes. **(A)** The CTD-CCD interface observed in cocrystals of BI-D-IN complexes adapted from (68). **(B)** HEK293T cells were transfected with WT proviral HIV-1_NL4- 3_ expression plasmid or a derivative carrying *pol* mutations for Y226A, W235A, K266A & I268A substitutions. Cell lysates and cell culture supernatants containing virions were collected 2 days post transfection and analyzed by immunoblotting for CA and IN. The image is representative of two independent experiments. **(C)** HEK293T cells were transfected as in panel B, and cell culture supernatants containing viruses were titered on TZM-bl indicator cells, whereby virus replication was limited to a single cycle by addition of dextran sulfate (50 μg/mL). The titers were normalized relative to particle numbers as assessed by an RT activity assay and are presented relative to WT (set to 1). Data show the mean from three independent replicates, error bars show the SD. Autoradiogram of IN-RNA adducts immunoprecipitated from virions bearing the indicated substitutions in IN. Immunoblots below show immunoprecipitated IN or IN protein in lysates prior to immunoprecipitation. Data shown are representative of three independent experiments. **(D)** Quantitation of IN CLIP autoradiographs. Data show the mean from three independent replicates, error bars show the SD. **(F-G)** HEK293T cells were transfected with full-length pNL4- 3-derived proviral plasmids carrying pol mutations coding for IN Y226A, W235A, K266A & I268A substitutions introduced on the IN-D278R backbone. Cell lysates and cell culture supernatants were analyzed as by immunoblotting for CA and IN **(F),** and cell culture supernatants containing viruses were titered on TZM-bl indicator cells **(G)** as above. **(H)** Autoradiogram of IN-RNA adducts immunoprecipitated from virions bearing the indicated substitutions in IN introduced on the IN- D278R backbone as in panel F. Immunoblots below show immunoprecipitated (IP) IN or CA protein in lysates prior to immunoprecipitation**. (I)** Quantitation of IN CLIP autoradiographs. Data show the mean from three independent replicates, error bars show the SD. Panels C, E, G, I, \*\*\**P* < 0.001; \*\*\*\**P* < 0.0001 by one-way ANOVA with Dunnett’s correction for multiple comparisons.

We previously demonstrated that the charge reversal substitution IN D278R restores infectivity to the IN R269A/K273A class II mutant by re-establishing IN binding to gRNA, highlighting the electrostatic nature of IN-gRNA interactions (60). We investigated whether the IN D278R substitution could similarly help suppress the replication defects of HIV-1_(Y226A IN)_, HIV-1_(W235A IN)_, HIV-1_(K266A IN)_ and HIV-1_(I268A IN)_. Like above, combined substitutions did not affect cell-associated Gag/Gag-Pol expression, Gag/Gag-Pol processing, particle release or IN levels in virus particles (**Fig. 1F**). However, the resultant viruses still exhibited significantly reduced viral infectivity (**Fig. 1G**), suggesting that Y226A, W235A, K266A and I268A mutations inhibit IN-gRNA binding through a different mechanism than the IN R269A/K273A mutant. On the other hand, we found that the IN D278R substitution helped stabilize IN and resulted in similar amounts of IP’ed IN protein (**Fig. 1H**). Notwithstanding, all mutants displayed class II phenotypes with IN–gRNA binding either reduced or abolished (**Fig. 1H**). Collectively, these results indicate that mutations within CTD residues involved in ALLINI-induced aberrant IN multimerization result in non-infectious viruses with the class II phenotype.

### HIV-1_(R269A/K273A/D278R IN)_ displays enhanced sensitivity to quinoline- and pyrrolopyridine- based ALLINIs

IN R269A/K273A substitutions within the CTD directly inhibit gRNA binding and result in the formation of morphologically aberrant eccentric particles that are non-infectious (20, 58, 60). The replication defect of HIV-1_(R269A/K273A IN)_ is rescued by restoring the local positive charge at nearby residues (e.g. D256R, D256N/D270N(D2N), D278R, and D279R (60)). Given their vicinity to the ALLINI interacting CTD residues (**Fig. 1A**), we reasoned that IN R269A/K273A substitutions may affect ALLINI sensitivity. To test this hypothesis, we first examined the ALLINI sensitivity of the HIV-1_(R269A/K273A/D278R IN)_ mutant virus.

Surprisingly, we found that the HIV-1_(R269A/K273A/D278R IN)_ was substantially (∼100-fold) more sensitive to quinoline-based ALLINI-2 than WT HIV-1 (IC_50_ for the mutant virus vs WT HIV-1: ∼3 nM vs ∼308 nM, **Fig. 2A**). This enhanced sensitivity correlated with reduced RNA binding by IN R269A/K273A/D278R at ALLINI-2 concentrations as low as 0.01 µM, whereas WT IN-RNA binding was unaffected at 0.01 and 0.1 µM (**Fig. 2B, 2C**). HIV-1_(R269A/K273A/D278R IN)_ was similarly more sensitive to another quinolone-based ALLINI, BI-D (IC_50_ for WT vs. the mutant virus vs WT HIV-1: ∼4 nM vs ∼376 nM, **Fig. 2D**) as well as pyrrolopyridine-based Pirmitegravir (IC_50_ for the mutant virus vs WT HIV-1: ∼0.09 nM vs. 29 nM, **Fig. 2G**). In both cases, increased sensitivity of the IN mutants correlated with loss of RNA binding at low concentrations of ALLINIs (**Fig. 2E, 2F, 2H, 2I**). These results demonstrate that the CTD substitutions away from the primary ALLINI- binding interface at the CCD dimer can alter sensitivity to different classes of ALLINIs.

**Figure 2.**
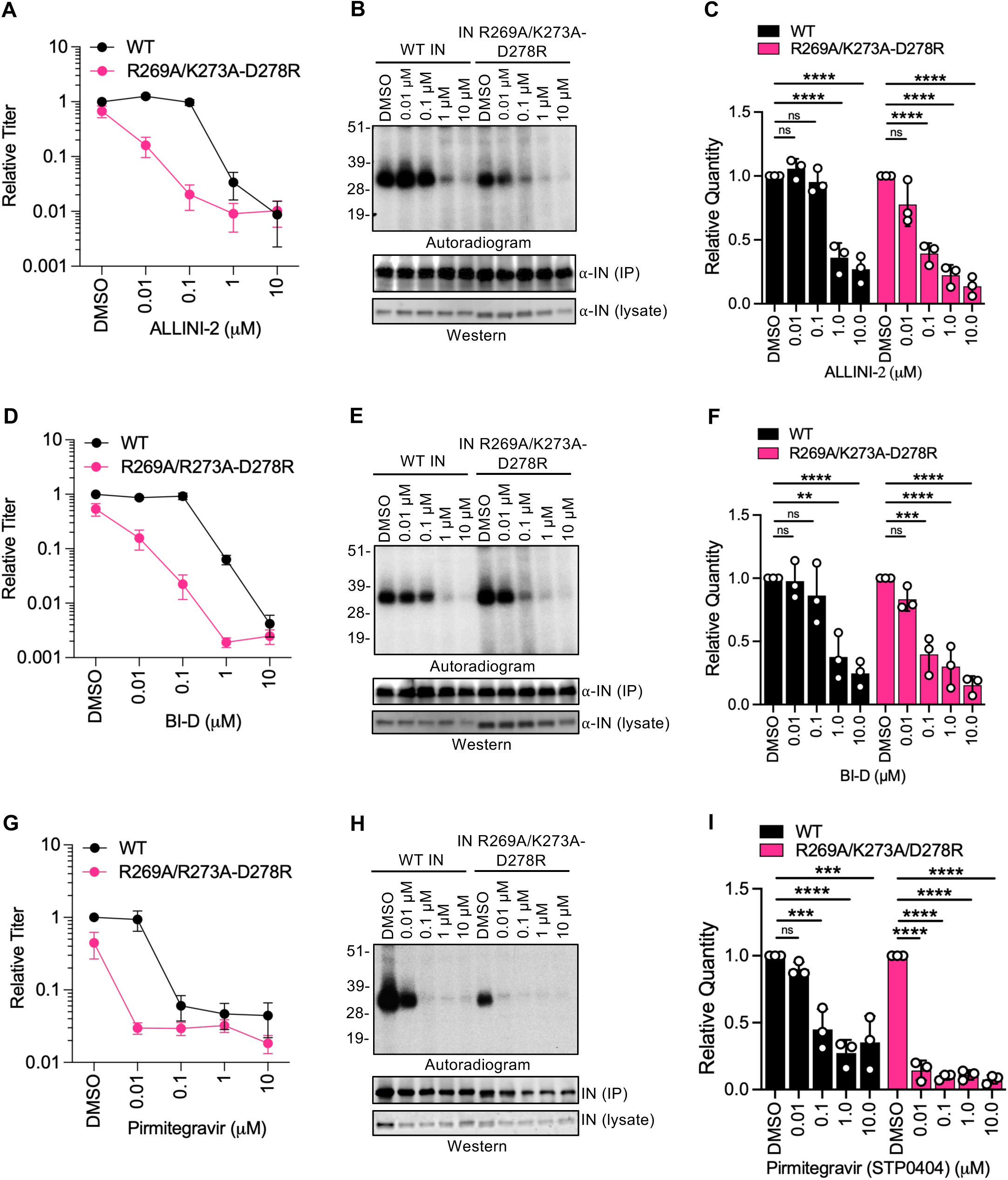
**IN R269A/K273A/D278R substitutions confer enhanced sensitivity to ALLINIs**. **(A- I)** HEK293T cells were transfected with WT proviral HIV-1_NL4-3_ expression plasmid or a derivative carrying *pol* mutations for IN R269A/K273A/D278R substitutions. Cells were subsequently treated with 0.01, 0.1, 1.0 & 10.0 µM of ALLINI-2 (A-C), BI-D (D-F) and Pirmitegravir (STP0404) (G-I). (**A, D, G**) Cell culture supernatants containing viruses were titered on TZM-bl indicator cells, whereby virus replication was limited to a single cycle by addition of dextran sulfate (50 μg/mL). The titers were normalized relative to particle numbers as assessed by an RT activity assay and are presented relative to WT. **(B, E, H)** Autoradiogram of IN-RNA adducts immunoprecipitated from virus particles harvested from the supernatants of ALLINI-treated cells. Immunoblots below show IN protein levels in lysates prior to immunoprecipitation and IP’ed IN. Data shown are representative of three independent experiments. **(C, F, I)** Quantitation of IN-RNA adducts from autoradiographs. IN-RNA complex band intensities were normalized relative to WT (set to 1). Data show the mean from three independent experiments, error bars display the SD. Ns (not significant), \*\**P* < 0.01, \*\*\**P* < 0.001; \*\*\*\**P* < 0.0001 by one-way ANOVA with Sidak’s correction for multiple comparisons.

### Enhanced ALLINI sensitivity of HIV-1_(R269A/K273A/D278R IN)_ maps to the R269A and K273A substitutions

We next examined whether the IN R269A/K273A class II substitutions or the IN D278R compensatory substitution contributed to the enhanced sensitivity of HIV-1_(R269A/K273A/D278R IN)_ to ALLINIs. To this end, we determined the ALLINI sensitivity of HIV-1_(R269A/K273A IN)_ bearing other compensatory substitutions (e.g. D256R, D256N/D270N-D2N, and D256R/D270R-D2R). IN_R269A/K273A_ bearing all three compensatory substitutions displayed similarly enhanced sensitivity to 0.1 µM ALLINI-2 (**Fig. 3A**) and BI-D (**Fig. 3B**), suggesting that the R269A/K273A class II substitutions but not the compensatory substitutions confer ALLINI sensitivity. This correlated with reduced IN-RNA binding at 0.1 µM BI-D, whereby WT IN was not affected (**Fig. 3C**). Another class II IN mutant, IN R262A/R263A bearing the compensatory D256R substitution was less sensitive to 0.1 µM ALLINI-2 and BI-D (**Fig. 3A, 3B**) and maintained RNA binding at 0.1 µM BI- D **(Fig. 3D**). Together, these findings suggest a role of the IN R269A/K273A substitutions in the observed enhanced sensitivity to ALLINIs.

**Figure 3.**
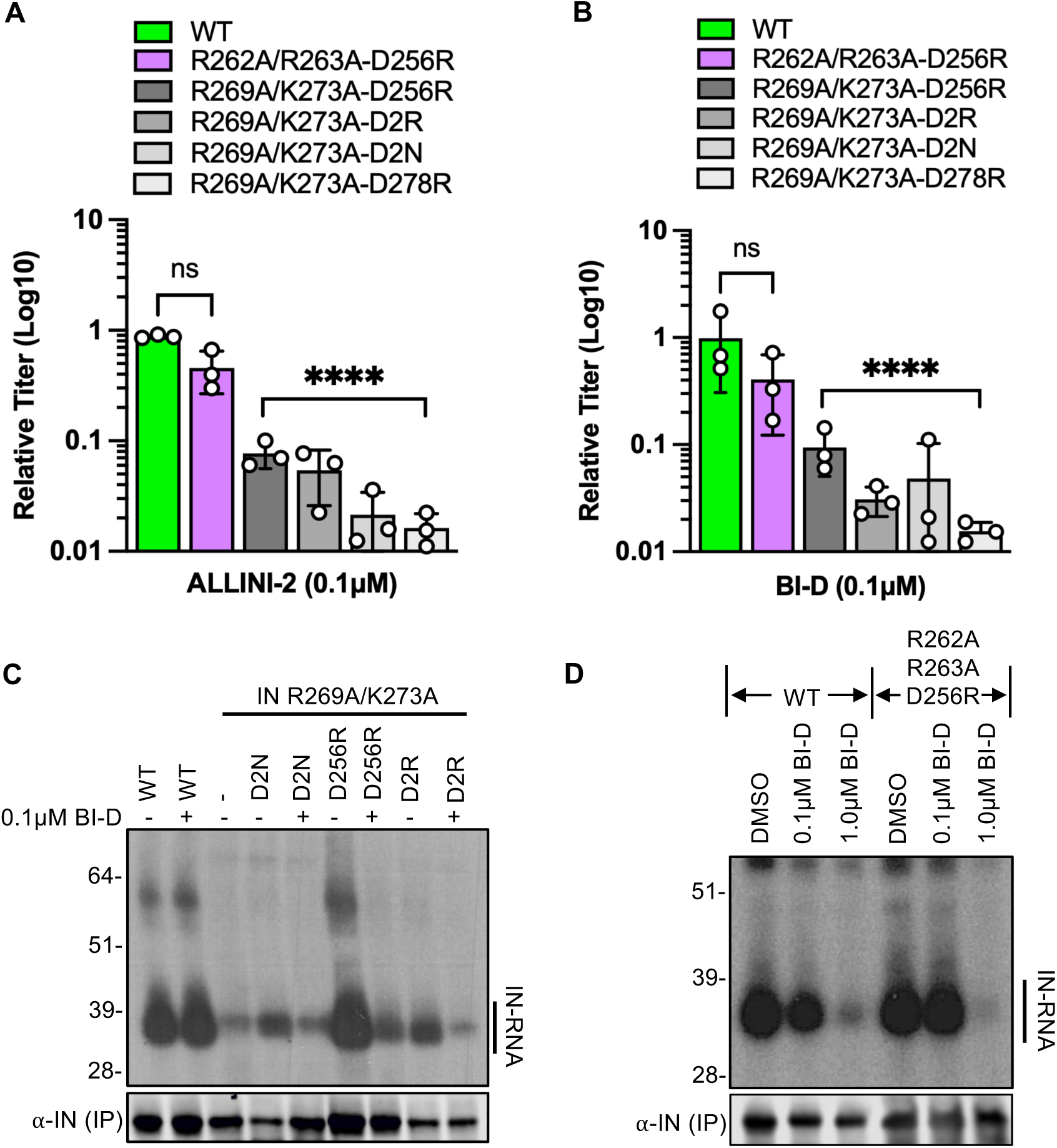
Enhanced ALLINI sensitivity of the IN R269A/K273A/D278R mutant maps to A269 and A273 residues. (A-D) HEK293T cells were transfected with proviral HIV-1_NL4-3_ expression plasmids carrying *pol* mutations for the indicated compensatory IN substitutions D256R, D2R and D2N in the background of IN R269A/K273A mutant, D256R compensatory substitution on IN R262A/R263A backbone, or the WT control. Cells were treated with 0.1 µM ALLINI-2 (A) or BI-D (B-D). Cell culture supernatants containing viruses were titered on TZM-bl indicator cells, whereby virus replication was limited to a single cycle by addition of dextran sulfate (50 μg/mL). The titers were normalized relative to respective mock-treated samples for each virus (set to 1) (A-B). Data show the mean from three independent experiments, error bars display the SD. Ns (not significant), \*\*\*\**P* < 0.0001 by one-way ANOVA with Dunnet’s correction for multiple comparisons. Autoradiogram of IN-RNA adducts immunoprecipitated from virions bearing the indicated IN substitutions (C, D). Immunoblots below show immunoprecipitated (IP) IN protein. Data shown are representative of three independent experiments.

### IN R269A/K273A/D278R substitutions do not enhance ALLINI-induced aberrant IN oligomerization

As ALLINIs lead to aberrant IN multimerization (62–67, 69), we examined whether IN R269A/K273A/D278R substitutions enhance ALLINI sensitivity due to more efficient multimerization at low ALLINI concentrations. Analysis of IN oligomerization using EGS-based protein-protein crosslinking revealed no major differences in oligomeric states between WT and R269A/K273A/D278R IN at low ALLINI concentrations (**Fig. 4A-D**). Furthermore, dynamic light scattering (DLS) analysis did not support increased susceptibility to aggregation for the IN R269A/K273A/D278R mutant compared to WT IN in the presence of ALLINI-2 (**Fig. 4E-H**). On the contrary, the IN R269A/K273A-D278R mutant appeared to form larger aggregates albeit at a slower rate than WT IN. Collectively, these data underscore the importance of IN–gRNA binding, but not enhanced aberrant IN multimerization, as a critical factor that underlies the increased ALLINI sensitivity of the IN R269A/K273A/D278R mutant.

**Figure 4.**
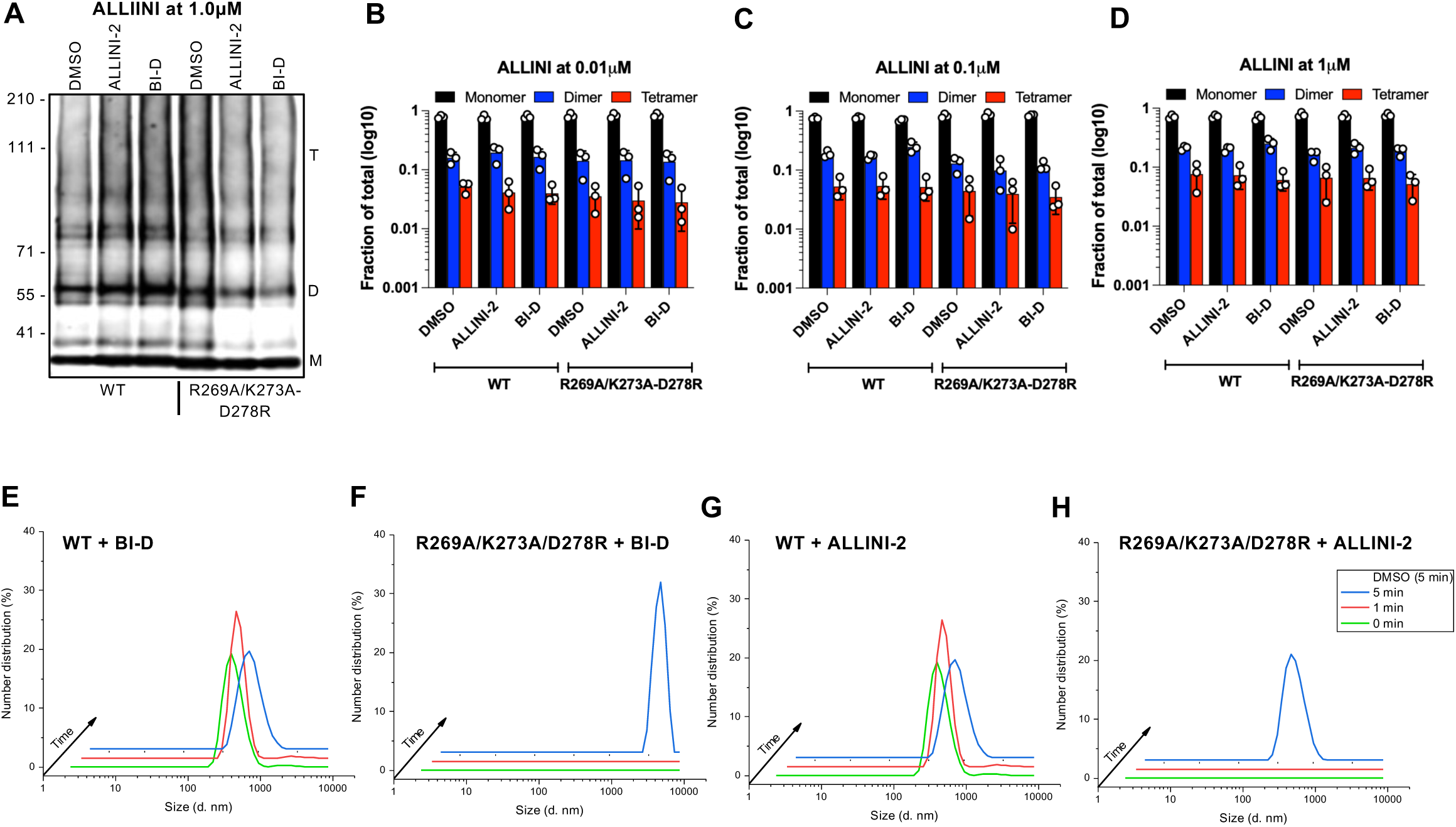
IN R269A/K273A/D278R mutant is not more prone to ALLINI-induced multimerization. (A-D) HIV-1_WT_ or HIV-1_(R269A/K273A/D278R IN)_ virus particles generated in the presence of 1.0 µM (A, B), 0.1 µM (C) and 0.01 µM (D) ALLINI-2 and BI-D were treated with 1 mM EGS for IN crosslinking *in virio*. Virus lysates were subsequently analyzed by immunoblotting using antibodies against IN following separation on 3–6% Tris-acetate gels as detailed in Materials and Methods. The position of monomers (M), dimers (D), and tetramers (T) are indicated in a representative western blot (A). Quantification of IN multimerization from independent western blots (B, C, D). Data show the mean, error bars display the standard deviation from three independent experiments. **(E-H)** 10 nM of recombinant WT and R269A/K273A/D278R IN proteins were mixed with 50 nM of ALLINI-2 or BI-D. DLS was performed at the indicated time points to monitor the distribution of the particle size (diameter in nm). Data are representative of 2 independent experiments.

### Compensatory A268V and A268T/D270N IN substitutions suppress the replication defect of HIV-1_(I268A IN)_

To directly investigate the contribution of ALLINI-interacting CTD residues in ALLINI mechanism of action, we set out to identify secondary site suppressors of IN Y226A, W235A, K266A and I268A substitutions through serial passaging experiments. Passaging experiments were set up in three replicate sets to maximize the chances of recovering compensatory substitutions. In line with the infectivity defects in single-cycle infection (**Fig. 1**), all mutant viruses displayed delayed replication kinetics (**Fig. 5A-C**). While the IN I268A mutant outgrew in each replicate experiment, Y226A and W235A outgrew only in two and one replicate experiments, respectively (**Fig. 5A-C**). HIV-1_(K266A IN)_ was fully suppressed (**Fig. 5A-C**), indicating that K266 is a highly constrained site with minimal evolutionary tolerance. Outgrown viruses from passage 1 were inoculated on fresh cells and virus growth was similarly monitored. At the end of passage 2, IN Y226A, W235A and I268A mutants replicated at or near WT kinetics (**Fig. 5D-F**). Deep sequencing of virion-associated RNA revealed that Y226A and W235A mutants both reverted to WT upon passaging. On the other hand, A268V substitution was detected at the end of passage 1 and got further fixed in passage 2 in replicates 1 and 3 (**Fig. 5D, 5F**). Interestingly, in replicate 2, A268T substitution arose in passage 1 with an accompanying D270N substitution at a low frequency (4%) (**Fig. 5B**). At the end of passage 2, A268T/D270N substitutions were both fixed (**Fig. 5E**).

**Figure 5.**
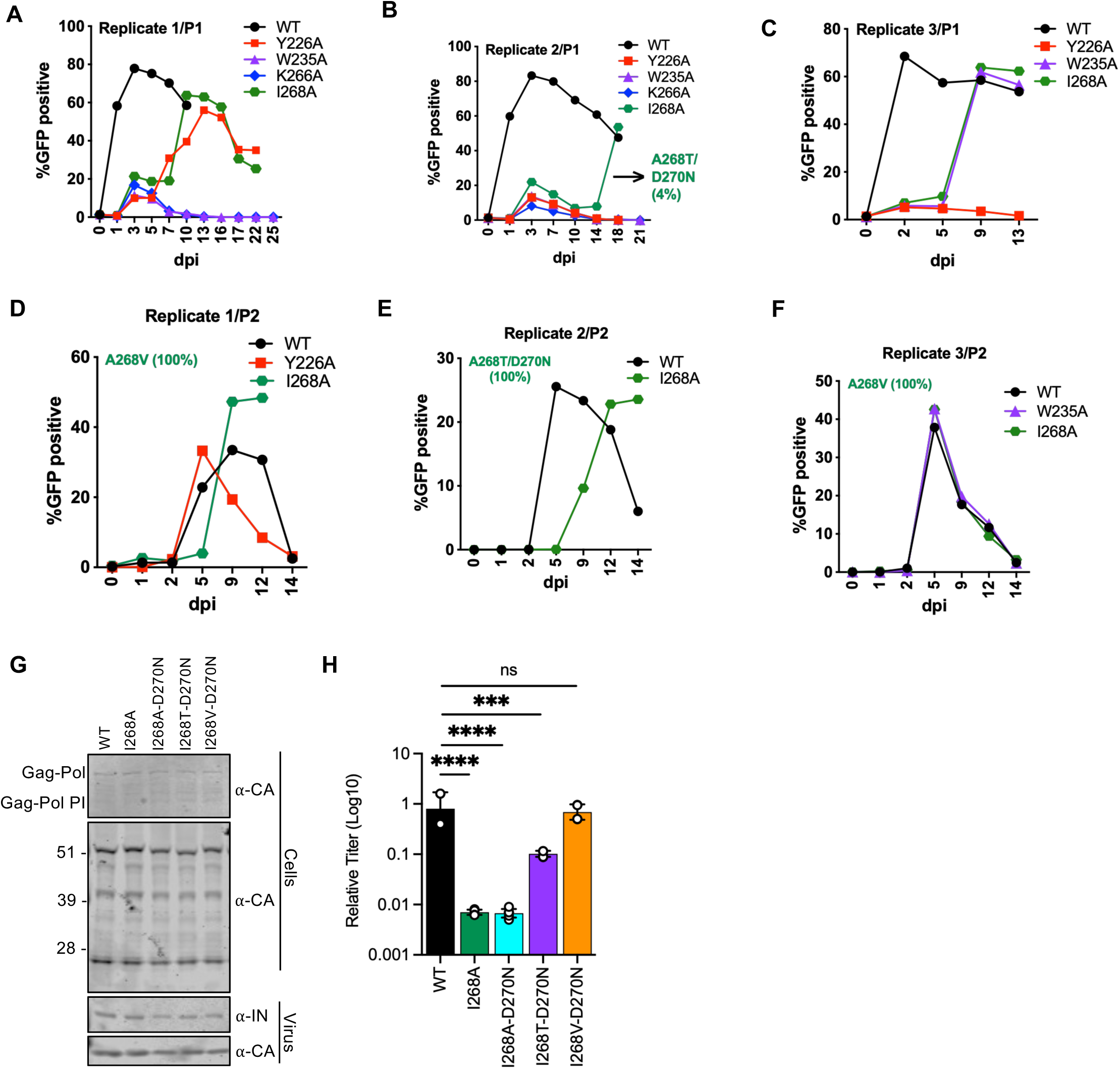
HIV-1 IN I268T/D270N and I268V compensatory substitutions suppress the replication defect of the HIV-1_(I268A IN)_ class II mutant virus. (A-F) MT4-LTR-GFP indicator cells were infected with WT HIV-1_NL4-3_ or derivatives bearing Y226A, W235A, K266A & I268A substitutions in IN in triplicate sets at an MOI of 2 as detailed in Materials and Methods. Viruses were serially passaged for two rounds until the emergence of compensatory mutations that allowed virus replication with WT kinetics. The graphs represent the percentage of GFP positive cells as assessed by flow cytometry over two passages (Panels A-C show passage 1, Panels D-F show passage 2) at the indicated days postinfection (dpi). HIV-1 genomic RNA was isolated from viruses collected from cell culture supernatants and analyzed by sequencing as described in Materials and Methods. Frequency of A268T/D270N and A268V substitutions in the sequencing data sets are indicated in figure insets. **(G, H)** HEK293T cells were transfected with full-length pNL4-3 expression plasmids carrying *pol* mutations coding for I268A, I268A-D270N, I268T- D270N and I268V-D270N substitutions or a WT control. Cell lysates and cell culture supernatants containing virus particles were collected 2 days post transfection and analyzed by immunoblotting for CA and IN (G). Cell culture supernatants containing viruses were titered on TZM-bl indicator cells, whereby virus replication was limited to a single cycle by addition of dextran sulfate (50 μg/mL). The titers were normalized relative to particle numbers as assessed by an RT activity assay and are represented relative to WT (H). Data show the mean from two independent experiments, error bars show the SD.

We next introduced the I268T/D270N and I268V/D270N substitutions in the NL4-3 proviral plasmid backbone and assessed viral gene expression, virion release and infectivity. We note that the I268V substitution did not arise together with the D270N substitution. Nonetheless, given that D270N substitution has no replicative cost on WT virus replication (60) and to better compare it to the IN I268T/D270N mutant, we decided to characterize the properties of the I268V/D270N double mutant in the following studies. The mutations had no impact on cell-associated Gag/Gag-Pol levels or processing, and particle release (**Fig. 5G**). While the IN I268V/D270N mutant was as infectious as WT, IN I268T/D270N mutant only partially restored infectivity (**Fig. 5H**). Of note, the compensatory effect was driven by substitutions at position 268, rather than D270N, as the I268A-D270N mutant remained noninfectious (**Fig. 5H**).

### IN A268T/D270N substitutions confer partial resistance to ALLINI-induced loss of gRNA binding

We next examined whether the IN I268T and I268V substitutions affected ALLINI sensitivity. HIV- 1_(I268V/D270N IN)_ displayed similar sensitivity to ALLINI-2 as WT HIV-1 (**Fig. 6A**), which correlated with reduced RNA binding at 1 µM (**Fig. 6B, 6C**). Despite being less infectious at baseline, HIV- 1_(I268T/D270N IN)_ maintained detectable levels of infectivity at 1 µM ALLINI-2 (**Fig. 6A**). Importantly, 1 µM ALLINI-2 which significantly reduced RNA binding by WT IN and IN I268V/D270N, decreased RNA binding by IN I268T/D270N only modestly and this reduction remained statistically nonsignificant (**Fig. 6B, 6C**). These findings were largely corroborated by similar results obtained with BI-D (**Fig. 6D-F**), though we noted that HIV-1_(I268V/D270N IN)_ was more sensitive to 0.1 µM BI-D as compared to WT HIV-1 (**Fig. 6D**). However, at this concentration, we didn’t observe a measurable reduction in RNA binding with IN I268V/D270N (**Fig. 6E, 6F**). Similar to ALLINI-2, 1 µM BI-D led to a significant reduction in RNA binding for both WT IN and IN I268V/D270N, whereas IN I268T/D270N remained more resistant to ALLINI-induced loss of RNA binding (**Fig. 6E, 6F**). These results collectively show that the Thr substitution at the ALLINI-interacting I268 residue confers resistance to ALLINI-induced loss of RNA binding, albeit this does not result in measurably enhanced viral titers.

**Figure 6.**
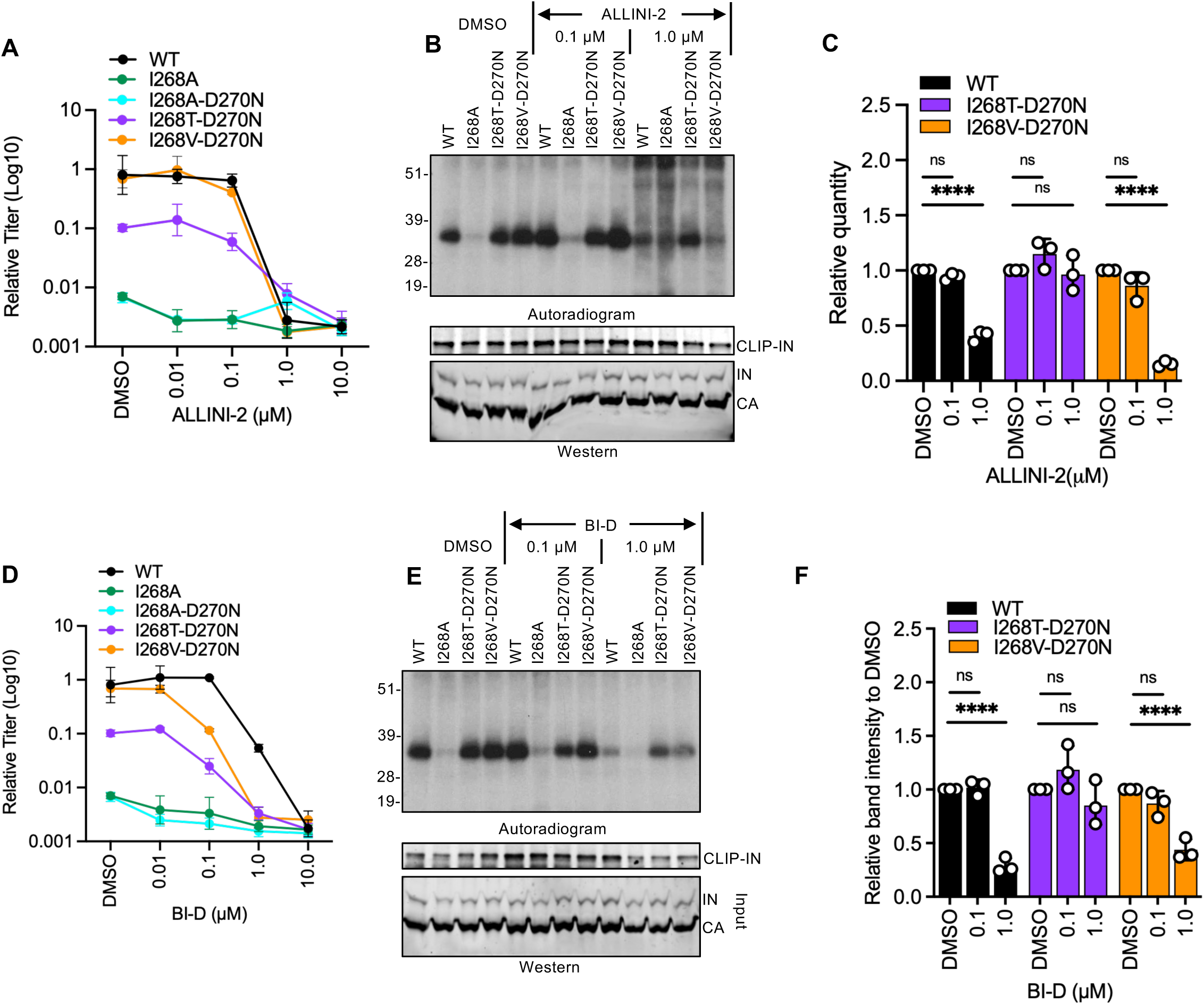
**Compensatory IN I268T/D270N substitutions confer resistance to ALLINI-induced block in RNA binding**. **(A, D)** HEK293T cells were transfected with proviral HIV-1_NL4-3_ expression plasmids carrying *pol* mutations for IN I268A, I268T-D270N and I268V-D270N, and treated with 0.01, 0.1, 1.0 or 10 µM of ALLINI-2 or BI-D. Virus titers were determined on TZM-bL cells and normalized relative to mock (DMSO)-treated WT infections. Data show the mean from 3 independent experiments, error bars show the SD. **(B, C, E, F)** HEK293T cells were transfected as above and treated with 0.1 or 1.0 µM of ALLINI-2 (B. C) or BI-D (D, E). Representative autoradiogram of IN-RNA adducts immunoprecipitated from virions (B, E). Immunoblots below show immunoprecipitated (IP) IN or CA protein in lysates prior to immunoprecipitation. Quantitation of IN-RNA adducts from autoradiographs (C, F). IN-RNA complex band intensities were normalized relative to mock-treated WT samples (set to 1). Data show the mean from three independent experiments, error bars display the SD. Ns (not significant), \*\**P* < 0.01, \*\*\**P* < 0.001; \*\*\*\**P* < 0.0001 by one-way ANOVA with Sidak’s correction for multiple comparisons.

## DISCUSSION

ALLINIs bind to a V-shaped pocket formed at the CCD dimer interface and recruit the CTD of an adjacent IN dimer. This aberrant multimerization through engagement of the IN-CTD, largely observed in biochemical studies, is thought to preclude IN binding to the gRNA in virions (61–69). The CTD has been central to this model, but previous studies failed to test its role within virions because substitutions of the CTD residues that contact the ALLINI-bound CCD yielded non- infectious virions. Indeed, here we confirm that alanine substitutions of Tyr226, Trp235, Lys266 and Ile268 give class II phenotypes with loss of IN-gRNA binding (**Fig. 1**). We have overcome this barrier by using compensatory CTD mutants that restore infectivity. Using this approach, we show that CTD substitutions lying outside the drug-binding pocket can either sensitize HIV-1 to ALLINIs (R269A/K273A) or partially protect IN-gRNA binding from ALLINI-induced disruption (I268T). To our knowledge, this provides the first virological evidence that the CTD contributes to ALLINI mechanism of action.

ALLINIs comprise of a bulky hydrophobic group (chromanyl and chlorophenyl in BI-D and PIR, respectively), a *tert*-butoxyl side chain, and an invariant carboxylate that are attached to an aromatic scaffold. Structural studies have shown that ALLINIs engage four invariant (Glu170, Tyr226, Lys266, and Trp235) and one highly conserved (Ile268) HIV-1 IN residues to facilitate CCD-CTD binding (68, 69). While the aromatic scaffolds of ALLINIs (e.g. quinoline in BI-D and pyrrolopyridine in PIR) engage in π-π stacking interactions with the side chains of Tyr226 and Trp235, Lys266 interacts with the side chain of CCD Glu170 and carboxylate groups of the inhibitors through ionic interactions. Within the CCD dimer interface, chromanyl and chlorophenyl groups of BI-D and PIR engage in hydrophobic interactions with the side chain of CTD Ile268. We predict that the introduction of the polar hydroxyl group by I268T substitution would weaken the hydrophobic contacts that help stabilize the ALLINI-induced CTD-CCD interface and allow IN remain engaged with the gRNA. The inability of the I268T/D270N mutant to restore infectivity likely suggests that these IN-gRNA complexes are not functional, likely due to the involvement of other residues involved in ALLINI-induced multimerization.

Hypersensitivity of the IN R269A/K273A/D278R to both quinoline- and pyrrolopyridine-based compounds indicates that sensitivity is mediated by the properties of CTD rather than chemotype-specific drug contacts. Hypersensitivity also tracked with the IN R269A/K273A substitutions rather than the identity of the compensatory substitution and was not a general feature of charge-compensated class II mutants, since R262A/R263A bearing the compensatory D256R substitution was not hypersensitive. The X-ray structure of BI-D bound to the CTD-CCD reveals that Lys273 is substantially distanced from the inhibitor binding pocket (68), whereas Arg269 is positioned immediately adjacent to Ile268 which directly contacts BI-D. The positively charged side chain of Arg269 points away from both CCD and BI-D to avoid the potential steric clashes and electrostatic repulsions. By contrast we predict that the R269A substitution introduces a small aliphatic side chain and thereby provide a more favorable hydrophobic environment for the CTD to effectively engage with the CCD + ALLINI complex.

A central question is whether the hypersensitivity of the R269A/K273A-bearing mutants arises from enhanced ALLINI-induced aberrant IN multimerization or from weakened IN-gRNA binding. Our data favor the latter. EGS crosslinking in virions showed no major differences in IN oligomeric states between WT HIV-1 and HIV-1_(R269A/K273A/D278R IN)_ at low ALLINI concentrations, and in DLS experiments the IN R269A/K273A/D278R mutant formed larger aggregates but did so more slowly than WT IN (**Fig. 4**). Therefore, we propose that R269A/K273A/D278R IN is more sensitive than WT IN to ALLINI-induced conformational changes that affect IN-RNA binding. The compensatory charge reversal restores infectivity, but the resulting IN-gRNA interface in the mutant virus could be more fragile and accordingly, more readily affected by ALLINIs than WT IN. This interpretation emphasizes IN-gRNA binding, rather than multimerization *per se*, as the functionally relevant target of ALLINIs. One limitation in our study is that DLS reports the kinetics of aggregate formation but not what fraction of soluble IN is converted to aggregates, and in-virion crosslinking provides a coarse readout of multimeric states of IN. More sensitive biophysical measurements will be needed to better delineate the contributions of ALLINI induced aberrant IN multimerization vs. ALLINI mediated local conformational changes in IN CTD that lead to inhibition of IN-vRNA binding.

ALLINI resistance substitutions selected in culture (Y99H, A128T, H171Q, T174I) cluster near the inhibitor-binding site in the CCD, where residues are variable, whereas none have been recovered in the CTD. The CTD residues that engage the ALLINI-CCD complex are invariant or highly conserved (67, 68), and they are essential for IN-gRNA binding and virion maturation. The one residue where adaptation was possible, Ile268, is the position where Leu occasionally occurs in circulating strains. Serial passaging of the four ALLINI-contacting CTD mutants supported these observations; Y226A and W235A reverted to WT, K266A failed to yield a replicating virus, and only I268A acquired true second-site substitutions, with a conservative change to a Val in one case. In the second case, it is noteworthy that the IN I268T/D270N is less infectious than WT at baseline and ALLINI-resistance conferred by I268T is also partial at best. Hence, it is unlikely that I268T could emerge as a clinical resistance route to ALLINIs. Given the high genetic barrier to CTD-mediated ALLINI resistance, our approach to use compensatory substitutions to restore infectivity to viruses bearing mutations in other ALLINI-interacting residues will be an important general strategy to map the critical ALLINI-CTD interactions involved in ALLINI mechanism of action moving forward.

Our findings suggest that natural CTD polymorphisms, or other perturbations that weaken IN-gRNA binding without abolishing infectivity, could sensitize virus to ALLINIs, and that compounds which exploit the CTD-RNA interface may enhance antiviral potency. Several questions remain. First, ALLINIs are added to producer cells, so it is unclear whether they can dislodge IN-gRNA complexes that have already formed, or whether they act mainly by preventing the formation of functional IN-gRNA complexes during assembly and maturation. Second, the relationship between ALLINI-induced multimerization and IN-gRNA displacement has not been well studied in vitro, and reconstituted assays with defined RNA would help address whether ALLINI-induced aggregates compete directly with RNA. In summary, our findings show that CTD residues distant from the ALLINI-binding pocket modulate drug sensitivity by tuning IN-gRNA binding and multimerization propensity, and they identify IN-gRNA binding as a central target of ALLINIs. These insights not only advance our understanding of HIV-1 integrase biology but also have important implications for the design of next-generation ALLINIs that exploit vulnerabilities within the CTD to achieve enhanced antiviral potency and a higher barrier to resistance.

## MATERIALS AND METHODS

### Plasmids and Compounds

IN mutations were introduced into the HIV-1_NL4-3_ -derived full-length proviral plasmids bearing WT IN or IN D278R by overlap extension PCR. In brief, primers containing IN mutations were used in PCR reactions with forward (with EcoRI restriction endonuclease site) and reverse (with AgeI restriction endonuclease site) outer primers. The resulting fragments containing the desired mutations were mixed in an overlap extension PCR by using the outer primer pairs. The overlap fragments were digested with AgeI and EcoRI and cloned into pNL4-3 plasmids. The presence of the desired mutations and the absence of unwanted secondary changes was assessed by Sanger sequencing. Viruses bearing class II substitutions (IN R262A/R263A, IN R269A/K273A) as well as those bearing additional compensatory substitutions have been described before (20, 58, 60). ALLINIs, BI-D and BI-B2, were synthesized by Aris Pharmaceuticals using previously published protocols (70, 71). Pirmitegravir (HY-130000) was obtained from MedChemExpress. Compounds were dissolved in DMSO at a final concentration of 10 mM and stored in -80°C.

### Cell lines and viruses

HEK293T cells (ATCC CRL-11268) and TZM-bL cells were cultured in Dulbecco’s modified Eagle’s medium supplemented with 10% fetal bovine serum. Hela-derived TZM-bL cells that express CD4/CXCR4/CCR5 receptor/coreceptors and bear an HIV-1 LTR driven β-galactosidase reporter were obtained from the NIH AIDS Reagent Program. Cell lines are regularly inspected for mycoplasma contamination using MycoAlert mycoplasma detection kit (Lonza) and checked for being free of any other contaminations. MT4-LTR-GFP (MT4-GFP) indicator cells bearing an HIV-1 LTR driven GFP reporter have been described before (72) and maintained in RPMI 1640 medium supplemented with 10% fetal bovine serum.

HEK293T cells grown in 24-wells plates were transfected with pNL4-3-derived plasmids bearing the abovementioned substitutions using polyethyleneimine (PolySciences). Media was replaced at 6 h post-transfection supplemented with different concentrations of ALLINIs as indicated in figure legends. At 2 days post-transfection, cell culture supernatants containing virions were titered on TZM-bL cells using a β-galactosidase assay (Galactostar, Thermo Fisher) per manufacturer’s instructions. Virus replication in TZM-bL cells was limited to a single-cycle via addition of dextran sulfate (50 mg/mL) 6-14 hours post-infection. An aliquot of the viruses were also subjected to a qPCR-based reverse-transcriptase (RT) activity assay (73). Virus titers obtained from infection of TZM-bL cells was normalized to particle numbers based on values obtained from the RT activity assays to determine the infectiousness (i.e. infectious units/particle) of viruses. A fraction of the cell culture supernatant collected from transfected HEK293T cells was also concentrated by centrifugation (16,100 x g, 4°C, 90min) on a 20% sucrose cushion prepared in 1x phosphate-buffered saline (PBS) for analysis of virion-associated proteins by immunoblotting as described below. In parallel, cells were lysed in 1xRIPA buffer and lysates used in immunoblotting as described below.

### Immunoblotting

Virus and cell lysates were resuspended in 1X sodium dodecyl sulfate (SDS) sample buffer, separated by electrophoresis on Bolt 4-12% Bis-Tris Plus gels (Life Technologies) and transferred to nitrocellulose membranes (Hybond ECL, Amersham). The membranes were probed overnight at 4°C with a mouse monoclonal anti-HIV capsid antibody (183-H12-5C, NIH AIDS reagents) or a mouse monoclonal anti-HIV integrase antibody (74) in Odyssey Blocking Buffer (LI-COR). Membranes were probed with fluorophore-conjugated secondary antibodies (LI-COR) and scanned using a LI-COR Odyssey system. IN and CA levels in virions were quantified using Image Studio software (LI-COR).

### Analysis of IN-RNA interactions by CLIP

CLIP experiments were conducted as previously described (58, 75–78). In short, cells in 15-cm cell culture plates were transfected with 30 μg full-length proviral plasmid (pNL4-3) DNA or derivatives carrying the IN mutations. Cell culture media was changed one-day post-transfection supplemented with different concentrations of ALLINIs and 4-thiouridine (4SU). Cell culture supernatants containing virions were collected two days post-transfection, filtered through 0.22 μm filters and pelleted by ultracentrifugation through a 20% sucrose cushion prepared in 1X PBS using a Beckman SW 32-Ti rotor at 29,000rpm for 1.5 hrs at 4°C. The virus pellets were resuspended in 1XPBS and UV-crosslinked for two consecutive times at an energy setting of 500 mJ in a Boekel UV-crosslinking chamber equipped with UV368 nm bulbs. Following lysis in 1X RIPA buffer, IN-RNA complexes were immunoprecipitated using a mouse monoclonal anti-IN antibody (74). Bound RNA was end-labeled with γ-^32^P-ATP and T4 polynucleotide kinase. The isolated protein-RNA complexes were separated by SDS-PAGE, transferred to nitrocellulose membranes (Hybond ECL, Amersham), and exposed to autoradiography films to visualize IN-RNA complexes. If needed, following immunoblotting and quantitation of immunoprecipitated IN, equivalent amounts of immunoprecipitated IN were re-run to accurately assess the ability of IN variants to bind RNA. Lysates and immunoprecipitates were also analyzed by immunoblotting using the aforementioned mouse monoclonal antibodies against IN. Protein-RNA complex amounts were quantified from scanned autoradiography films using the Image Studio software (LI-COR).

### ALLINI-induced IN aggregation by Dynamic Light Scaterring (DLS)

The assays were performed on a Malvern Zetasizer Nano s90 as described previously (64). In brief, recombinant WT and mutant INs at 10 nM were mixed with 50 nM of either ALLINI-2 or BI-D. Kinetic analysis was carried out at specified time points. In short, the reactions were performed in DLS buffer (1M NaCl, 2 mM MgCl_2_, 2 mM DTT, and 50 mM HEPES pH 7.5) which was filtered twice using 0.2 µm filter units. Drugs were prepared in DMSO at 10 mM stock concentration. 0.2 μL of serially diluted drug was added to 40 μL IN and size distributions of the mixture were recorded at 1 min intervals (0, 1, 2, 3, 4, and 5 min) until it reaches a total time of 5 min incubation. For negative control, the same amount of IN was mixed with 0.2 µL DMSO in the absence of drug.

### Assessment of IN multimerization in virions

HEK293T cells grown in 15-cm dishes were transfected with 30 μg pNL4-3 plasmid DNA bearing WT IN or R269A/K273A-D278R IN. One day post-transfection, cell culture media was replaced with media supplemented with 0.01µM, 0.1µM and 1.0µM of ALLINIs. Two days post-transfection, cell culture supernatants containing virions were filtered through 0.22 μm filters and pelleted by ultracentrifugation through a 20% sucrose cushion prepared in 1X PBS using a Beckman SW 32- Ti rotor at 29,000rpm for 1.5 hrs at 4°C. Pelleted virions were resuspended in 1xPBS and treated with a membrane-permeable crosslinker, EGS (ThermoFisher Scientific), at a concentration of 1mM for 30 min at room temperature. Crosslinking was stopped by the addition of 4XSDS-PAGE sample buffer. The cross-linked samples were then separated on 3-8% Tris-acetate gels and analyzed by immunoblotting using a mouse monoclonal anti-IN antibody (74).

### Serial passaging experiments and identification of secondary site suppressors

VSV-G-pseudotyped HIV-1 NL4-3 bearing WT IN or IN Y226A, W235A, K266A and I268A mutations were generated by transfecting HEK293T cells with proviral plasmids alongside with a VSV-G expression plasmid at a 9:1 ratio using polyethyleneimine (PolySciences, Warrington, PA). Cell culture supernatants containing viruses were collected 2 days post-transfection, filtered, aliquoted, and stored at −80°C. Virus titers were determined on TZM-bL cells as above and relative particle numbers in each virus stock assessed by an RT activity assay (73). Two million MT4-GFP cells were inoculated at a multiplicity of infection (MOI) 1 for WT virus or an equivalent particle number of mutant viruses. Infected cells were split at a ratio of 1:5 every 3–4 days and infections were monitored by flow cytometry of GFP-positive cells at the indicated intervals shown in figure legends. Cell culture supernatants containing virions were also collected at the same time intervals for subsequent passaging and sequencing analyses. At the end of each passage, cell culture supernatants containing infectious virions were collected and titered on MT4-GFP cells and used to inoculate fresh MT4-GFP cells as above. Aliquots of infected cells and viral particles in the cell culture supernatants were collected over the duration of each passage as above.

For sequencing analysis of viruses bearing compensatory mutations, virions collected at peak replication time points were concentrated on 20% sucrose cushions prepared in 1X PBS by ultracentrifugation (28,000 rpm, 4°C, 90 min, Beckman SW41 rotor). Genomic RNA was isolated from pelleted virions using TRIzol per manufacturer’s instructions. Extracted RNA was prepared for deep sequencing using the Illumina TruSeq Stranded Total RNA library kit following the manufacturer’s instructions but omitting the rRNA depletion step. Resulting libraries were sequenced on an Illumina HiSeq 2000 platform at the Genome Technology Access Center at Washington University School of Medicine. Sequencing reads were mapped to the pNL4-3 reference viral genome allowing for 2 mismatches using the Bowtie algorithm (i.e., -v 2, -m 10 parameters) (96), and the frequency of compensatory mutations was determined thereafter.

## ACKNOWLEDGEMENTS

This work was supported by NIH grants AI150497 (SBK), AI1508470 (U54 Center for HIV RNA Studies, SBK, RG), R01 AI143649 (MK) and Milton Schlesinger Student Fellowship (CSM, AB).

